# Spatiotemporal patterns in cortical development: Age, puberty, and individual variability from 9 to 13 years of age

**DOI:** 10.1101/2024.06.29.601354

**Authors:** Katherine L. Bottenhorn, Jordan D. Corbett, Hedyeh Ahmadi, Megan M. Herting

**Affiliations:** Department of Population and Public Health Sciences, University of Southern California, Los Angeles, CA, USA; Department of Psychology, Florida International University, Miami, FL, USA

**Keywords:** brain development, brain function, adolescent, child, puberty

## Abstract

Humans and nonhuman primate studies suggest that timing and tempo of cortical development varies neuroanatomically along a sensorimotor-to-association (S-A) axis. Prior human studies have reported a principal S-A axis across various modalities, but largely rely on cross-sectional samples with wide age-ranges. Here, we investigate developmental changes and individual variability in cortical organization along the S-A axis between the ages of 9-13 years using a large, longitudinal sample (N = 2487-3747, 46-50% female) from the Adolescent Brain Cognitive Development Study (ABCD Study®). This work assesses multiple aspects of neurodevelopment indexed by changes in cortical thickness, cortical microarchitecture, and resting-state functional fluctuations. First, we evaluated S-A organization in age-related changes and, then, computed individual-level S-A alignment in brain changes and assessing differences therein due to age, sex, and puberty. Varying degrees of linear and quadratic age-related brain changes were identified along the S-A axis. Yet, these patterns of cortical development were overshadowed by considerable individual variability in S-A alignment. Even within individuals, there was little correspondence between S-A patterning across the different aspects of neurodevelopment investigated (i.e., cortical morphology, microarchitecture, function). Some of the individual variation in developmental patterning of cortical morphology and microarchitecture was explained by age, sex, and pubertal development. Altogether, this work contextualizes prior findings that regional age differences do progress along an S-A axis at a group level, while highlighting broad variation in developmental change between individuals and between aspects of cortical development, in part due to sex and puberty.

**Significance Statement:** Understanding normative patterns of adolescent brain change, and individual variability therein, is crucial for disentangling healthy and abnormal development. We used longitudinal human neuroimaging data to study several aspects of neurodevelopment during early adolescence and assessed their organization along a sensorimotor-to-association (S-A) axis across the cerebral cortex. Age differences in brain changes were linear and curvilinear along this S-A axis. However, individual-level sensorimotor-association alignment varied considerably, driven in part by differences in age, sex, and pubertal development.

## Introduction

Human brain development begins *in utero* and extends through childhood and adolescence, into early adulthood (Sowell et al., 2004; Wierenga et al., 2014). During adolescence, synapse refinement, myelination, and apoptosis underlie functional plasticity and the maturation of large-scale neural circuitry (Huttenlocher, 1979; Schalbetter et al., 2022). The timing and pace of such development proceeds heterogeneously across brain regions. For example, synaptic pruning extends into late childhood in humans, ending years earlier in primary sensory regions than in prefrontal cortex (Huttenlocher and Dabholkar, 1997). Human magnetic resonance imaging (MRI) research echoes these neuronal findings, showing earlier maturation of gray matter morphology in unimodal somatosensory and visual cortices, but later maturation of higher-order association cortex (Gogtay et al., 2004). Several structural, functional, genetic, and metabolic features of the brain are similarly organized, with a hierarchical topology, indicating a sensorimotor-association (S-A) axis of cortical organization (Margulies et al., 2016; Sydnor et al., 2021). Further, some suggest that developmental plasticity across childhood and adolescence progresses along the cortical mantle such that unimodal, sensorimotor regions (e.g., visual cortex) mature prior to more heteromodal, associative regions (e.g., prefrontal cortex) (Gogtay et al., 2004; Dong et al., 2021; Sydnor et al., 2023). However, extant literature on S-A patterning of childhood and adolescent neurodevelopment has a few key shortcomings. First, puberty is infrequently considered in this work, although it is a critical neuroendocrine phenomenon of this developmental period featuring dramatic increases in hormones with central nervous system actions. While some work suggests that puberty does not play a role in S-A patterning beyond that of age (Sydnor et al., 2023), others find that puberty bisects peaks in sensorimotor and association regions’ cortical development (Grydeland et al., 2019). Further, S-to-A patterns have been uncovered in a number of neural phenomena, but no work to date has presented a multimodal assessment of S-to-A neurodevelopmental patterning. Here, we address these limitations by assessing S-A patterning in several aspects of cortical development during the transition to adolescence–and the roles of age, sex, and puberty therein–using multimodal MRI data from a large, longitudinal cohort.

We estimated individual-level developmental changes during the transition to adolescence using two waves of neuroimaging data from the Adolescent Brain Cognitive Development^SM^ Study (ABCD Study®), collected at ages 9-10 and 11-13 years, using multiple MRI phenotypes. *Cortical thickness* measurements approximate macroscale changes in cortical morphology due to synaptic pruning and myelination (Huttenlocher and Dabholkar, 1997; Asan et al., 2021). *Intracellular diffusion* from restriction spectrum imaging (RSI) approximates changes in cortical microarchitecture due to myelination, dendritic arborization, and apoptosis (White et al., 2012). Finally, *functional fluctuations*, estimated from the magnitude of intrinsic low frequency blood-oxygen level-dependent (BOLD) signal oscillations, approximates neural plasticity (Tolonen et al., 2007; Newbold et al., 2020; Laumann and Snyder, 2021). First, we assessed patterns of age-related brain changes along the S-A axis (Sydnor et al., 2021). We hypothesized that measures generally increasing across this developmental period of ages 9-13 years (i.e., isotropic intracellular diffusion) would exhibit greater increases over time in association areas than in the sensorimotor regions that are thought to develop earlier. In contrast, we expected measures generally decreasing across this developmental period (i.e., cortical thickness, directional intracellular diffusion) would show greater decreases over time in associative than sensorimotor regions. Finally, in measures that both increase and decrease in magnitude between ages 9-13 years (i.e., functional fluctuations), we expected to see increases over time in association areas and decreases, in sensorimotor areas. Then, we more thoroughly investigated potential age-, sex-, and pubertal-related individual differences in S-to-A development. Exploring the S-A alignment of longitudinal changes between ages 9-13 years in cortical morphology, microarchitecture, and function provides foundational knowledge necessary for neuroanatomical theories of cortical development. Furthermore, understanding normative patterns of adolescent brain change, and the extent of individual variability therein, is crucial for furthering multimodal MRI’s utility for disentangling healthy and abnormal development and for identifying potential intervention targets.

## Methods

### Participants

Data used here are from theAdolescent Brain Cognitive Development^SM^ Study (ABCD Study®) included in the 5.1 annual data release (http://dx.doi.org/10.15154/z563-zd24). The ABCD Study is a 10-year longitudinal, cohort study that recruited and enrolled 11,880 children 9 to 10 years of age at 21 study sites across the United States. Inclusion criteria were: English proficiency (of youth), in addition to absence of MRI contraindications or severe sensory, neurological, medical, or intellectual limitations (Garavan et al., 2018). The ABCD Study cohort closely matches the distribution of age, sex, and household size of 9- and 10-year-olds in the American Community Survey (ACS), a large, annual probability sample survey of U.S. households done by the U.S. Bureau of Census (Heeringa & Berglund, 2020). All children provided written, informed assent and their legal guardian gave informed consent and permission for the child to participate, per procedures approved by the centralized institutional review board and human research protections programs at the University of California San Diego. See Garavan et al. (Garavan et al., 2018) and Volkow et al. (Volkow et al., 2018) for more information. From the ABCD Study dataset, the current project utilized MRI data from the first two time points (i.e., baseline data collection and a follow-up visit two years later), in addition to participants’ sex at birth, age, and sociodemographic information from baseline data collection. Pubertal status estimates were obtained from the first three time points (i.e., baseline data collection (ages 9-10 years), 1-year follow up (ages 10-11 years), and 2-year follow up (ages 11-13 years)). Here, we further excluded data from individuals who had incidental findings on their structural MRI scans, whose MRI data failed the ABCD Study quality control procedures, and whose 2-year follow-up visit was on or after March 1, 2020, to mitigate any impact the COVID-19 pandemic may have on the findings. We further excluded imaging data from individuals exhibiting excessive head motion. For the current study, we also excluded data collected on General Electric (GE) and Philips Systems scanners due to substantial inter-scanner heterogeneity and software updates between data collection time points. Depending on MRI modality, our final analytic sample size ranged from 3474 to 2487 participants (**Table 1**).

**Table 1.** Participant Demographics.

|  | Cortical Thickness |  | Cortical Intracellular Diffusion |  | Resting-state Functional Fluctuations |  |
| --- | --- | --- | --- | --- | --- | --- |
|  | N | % of sample | N | % of sample | N | % of sample |
| Total N | 3474 | 30% | 2967 | 25% | 2487 | 21% |
| Age at baseline (months) | 119.50 ± 7.38 | | 119.59 ± 7.43 | | 119.93 ± 7.51 | |
| Sex (female) | 1596 | 46% | 1395 | 47% | 1241 | 50% |
| Puberty at baseline (Tanner) |  |  |  |  |  |  |
| Pre-pubertal | 1769 | 51% | 1540 | 52% | 1282 | 52% |
| Early puberty | 814 | 23% | 693 | 23% | 581 | 23% |
| Mid-puberty | 752 | 22% | 625 | 21% | 531 | 21% |
| Late/post puberty | 43 | 1% | 37 | 1% | 31 | 1% |
| Puberty change |  |  |  |  |  |  |
| Puberty | 0.40 ± 0.40 | | 0.40 ± 0.43 | | 0.42 ± 0.423 | |
| Puberty Missing | 429 | 12% | 343 | 12% | 272 | 0% |
| Race & Ethnicity |  |  |  |  |  |  |
| Asian + Other | 334 | 10% | 271 | 9% | 215 | 9% |
| Hispanic | 599 | 17% | 514 | 17% | 427 | 17% |
| Non-Hispanic Black | 465 | 13% | 354 | 12% | 258 | 10% |
| Non-Hispanic White | 2076 | 60% | 1828 | 62% | 1587 | 64% |
| Household Income |  |  |  |  |  |  |
| ≤ \$50,000 | 833 | 24% | 668 | 23% | 527 | 21% |
| \$50,000 to \$100,000 | 1035 | 30% | 889 | 30% | 750 | 30% |
| ≥ \$100,000 | 1365 | 39% | 1214 | 41% | 1053 | 42% |
| Caregiver Education |  |  |  |  |  |  |
| Up to high school diploma, GED | 111 | 3% | 80 | 3% | 55 | 2% |
| High school diploma, GED | 264 | 8% | 200 | 7% | 145 | 6% |
| Some college, Associate's degree | 924 | 27% | 769 | 26% | 631 | 25% |

**Table 1.** Participant Demographics
|  | Cortical Thickness |  | Cortical Intracellular Diffusion |  | Resting-state Functional Fluctuations |  |
| --- | --- | --- | --- | --- | --- | --- |
|  | N | % of sample | N | % of sample | N | % of sample |
| Bachelor's degree | 999 | 29% | 879 | 30% | 745 | 30% |
| Graduate degree | 1170 | 34% | 1033 | 35% | 908 | 37% |

*Note.* Within-category percentages that add to <100% are due to missing data and participants who declined to answer, i.e., in the case of household income. Abbreviations: ABCD, Adolescent Brain Cognitive Development; GED, General Educational Development test; HS, high school; MRI, magnetic resonance imaging; NA, not applicable. The “Other” race and ethnicity category includes participants who were identified by their parents as American Indian/Native American or Alaska Native; Asian Indian, Chinese, Filipino, Japanese, Korean, Vietnamese, or other Asian; Native Hawaiian, Guamanian, Samoan, or other Pacific Islander; or other race.

### Demographic and Developmental Data

Demographic data was reported by the participant’s parent or legal guardian (hereafter, “caregiver”) using the PhenX Toolkit (Hamilton et al., 2011). Age, in months, was calculated based on the participant’s birth date, as reported by their caregiver, and sex was based on their caregiver-reported sex assigned at birth. Caregivers reported what race and ethnicity they considered their child to be, which was then grouped into five categories for consistency with other socioeconomic data sources and constructs: Non-Hispanic White, Non-Hispanic Black, Hispanic, and Other (including Non-Hispanic Asian, American Indian, Native Hawaiian, Samoan, Chamorro, Other Pacific Islander, belonging to more than one race (e.g., multiracial) and other racial and ethnic identities). Household income was reported by the caregiver as the total annual income, from all sources in the past calendar year, which we summarized as ≤$50,000, $50,000 to $100,000, and ≥$100,000. Educational attainment was reported by the caregiver about their or their partner’s own highest level of education.

Pubertal status was calculated as a Tanner-equivalent category score based on the Pubertal Development Scale (Petersen et al., 1988), completed by the caregiver about the participant (Barch et al., 2018; Herting et al., 2021). The PDS consists of 5 questions, three of which are common to female and male youth and two of which differ by sex. The three common questions assess height growth, body hair, and skin changes. Questions for male youth assess vocal changes and facial hair, while questions for female youth assess breast development and menarche. For each of the 5 questions, parents/caregivers and youth were asked to separately rate their development on a 4-point scale (1 = has not begun yet, 2 = barely begun, 3 = definitely begun, 4 = seems complete), except for the menarche question for females, which consisted of a yes/no answer choice. Each question also consisted of an “I don’t know” answer. Here, we chose to use the caregiver-reported data due to a high number of “I don’t know” responses from youth at baseline data collection (i.e., between ages 9 and 10 years) (Cheng et al., 2021; Herting et al., 2021). A pubertal category score was derived for male participants by summing the body hair growth, voice change, and facial hair items and categorizing them as follows: prepuberty = 3; early puberty = 4 or 5 (no 3-point responses); midpuberty = 6-8 (no 4-point responses); late puberty = 9-11; postpuberty = 12. The puberty category score was derived for female participants by summing the body hair growth, breast development, and menarche and categorizing them as follows: prepubertal = 3; early puberty = 3 and no menarche; midpuberty = 4 and no menarche; late puberty ≤ 7 and menarche; postpuberty = 8 and menarche. Pubertal tempo was calculated using PDS stage estimates collected at ages 9-10 years (i.e., baseline data collection), ages 10-11 years, and ages 11-13 years by summing change between subsequent time points and dividing by the elapsed time.

### Neuroimaging Data

Data were acquired on SIEMENS 3T MRI scanners at each of 13 sites across the United States, using harmonized protocol and sequences, prior to which participants completed motion compliance training (Casey et al., 2018).

#### Cortical Thickness

T1w images were acquired using a magnetization-prepared rapid acquisition gradient echo (MPRAGE) sequence with integrated prospective motion correction, consisting of 176 slices with 1 mm^3^ isotropic resolution. The scanning protocol is further detailed by (Casey et al., 2018). Using Freesurfer (v7.1.1) cortical regions were labeled using the Desikan & Killiany atlas (Desikan et al., 2006) and average cortical thickness was computed per region of interest (ROI) (Fischl and Dale, 2000; Joyner et al., 2009; Rimol et al., 2010, 2010; Chen et al., 2012).

#### Resting State Functional Fluctuations

Four five-minute resting-state fMRI scans (eyes open), in two sets of two scans, were acquired using an echo-planar imaging sequence in the axial plane, with the following parameters: TR = 800 ms, TE = 30 ms, flip angle = 90°, voxel size = 2.4 mm^3^, 60 slices (Casey et al., 2018). Real-time head motion was monitored throughout the scans to reach the ABCD Study’s standards for low-motion data (>12.5 minutes of data). Here, we only include images without clinically significant incidental findings (*mrif_score* = 1 or 2) (Li et al., 2021) that passed all ABCD quality-control parameters were included in analysis (*imgincl_rsfmri_include* = 1), with at least 10 minutes of low-motion fMRI data (i.e., with framewise displacement (FD) < 0.2 mm) (Power et al. 2014).

Image processing is detailed by Hagler and colleagues (2019). Briefly, average preprocessed time series were extracted from each cortical ROI (in surface space). Further censoring was performed after time series were extracted from each ROI at FD > 0.2mm for network connectivity and BOLD variance estimates, removing high-motion volumes and time periods with fewer than five contiguous volumes (post-censoring). Then, outlier volumes were identified based on the standard deviation (SD) per ROI across TRs and time points with SD greater than three times the mean absolute deviation above or below the median SD were excluded. Functional fluctuations were calculated from each ROI, across the time series, and reflect the magnitude of low-frequency BOLD fluctuations.

#### Cortical Intracellular Diffusion

The DWI acquisition included a voxel size of 1.7 mm isotropic and with multiband acquisition (Moeller et al., 2010; Setsompop et al., 2012) with slice acceleration factor 3. Each DWI acquisition included a fieldmap scan for B0 distortion correction. ABCD employs a multi-shell diffusion acquisition protocol that includes 7 b=0 frames as well as 96 total diffusion directions at 4 b-values (6 with b = 500 s/mm^2^, 15 with b = 1000 s/mm^2^, 15 with b = 2000 s/mm^2^, and 60 with b = 3000 s/mm^2^). All images underwent distortion correction, bias field correction, motion correction, and manual and automated quality control per the steps detailed by Hagler and colleagues (2019). After preprocessing, white matter tracts were located and labeled according to the probabilistic atlas AtlasTrack (Hagler et al., 2009). Only images without clinically significant incidental findings (*mrif_score* = 1 or 2) that passed all ABCD quality-control parameters were included in analysis (*imgincl_dmri_include* = 1).

Restricted spectrum imaging (RSI) is a more advanced modeling technique that utilizes all 96 directions collected as part of ABCD’s multi-shell acquisition protocol. RSI provides detailed information regarding both the extracellular and intracellular compartments of white matter within the brain (White et al., 2013a, 2013b, & 2014). RSI model outputs include 7 normalized measures, all of which are unitless on a scale of 0 to 1. Restricted normalized isotropic signal fraction (RNI) measures *intracellular isotropic diffusion* and a higher RNI could indicate an increase in *cellularity*: the number of support cells or swelling of support cells, such as activated astrocytes or microglia (Palmer et al., 2022). Restricted normalized directional signal fraction (RND) measures *intracellular anisotropic diffusion* and likely indicates intra-axonal diffusion, with higher RND indicating greater *neurite density* from myelination and/or axonal packing (Palmer et al., 2022). Restricted normalized total signal fraction (RNT) indicates the total intracellular diffusion within a voxel and an increase could represent myelination, dendritic sprouting, arborization or increases in neurite density (Palmer et al., 2022). Mean RSI measures are calculated for cortical ROIs included in the Desikan and Killiany atlas (Desikan et al., 2006). For more details on image processing, see work by Hagler and colleagues (2019).

### Analyses

Prior to data analysis, a pre-registered analysis plan was registered with the [Open Science Framework (OSF)](https://doi.org/10.17605/OSF.IO/K3BYA). Significance thresholds for all reported models were adjusted for multiple comparisons using familywise error correction, adjusted for the number of effective comparisons between measures of cortical change (Li & Ji, 2005; Šidák 1969). Reported “significance” throughout the results refers to statistics with a *p*-value beneath this threshold. Analyses were performed using Python (v3.8.9) and R (v4.2.0) and all relevant code is available on [GitHub](https://github.com/62442katieb/deltaABCD_variability) (https://doi.org/10.5281/zenodo.12594702).

#### Missing data and multivariate outliers

From the entire ABCD Study baseline sample of 11,880 children, only 7457 (∼63%) children had imaging data from the 2-year follow-up appointment. From these, restricting our sample to individuals with high-quality MRI data at both time points (i.e., in order to calculate changes in MRI phenotypes), collected on SIEMENS scanners, yielded 3,474 (29% of baseline total, 46% female) children with high-quality structural MRI data, 2967 (25% of baseline total, 47% female) with high-quality diffusion-weighted MRI data, and 2487 (21% of baseline total, 50% female) with high-quality resting-state fMRI data.

Missingness was assessed, per variable, to determine (a) the proportion of missing values in the pre-COVID sample, (b) whether missingness was associated with values of other variables (i.e., missing at random), and (c) patterns of missingness across variables (Supplementary Methods, Supplementary Figure 1). Some participants have missing imaging data due to quality-related censoring (Supplementary Tables 1, 2). The presence of multivariate outliers in MRI phenotypes (i.e., both participant outliers and regional outliers) was assessed using random forests to recursively identify isolated points. No multivariate outliers were detected for cortical morphology, cortical microarchitecture, or resting-state functional fluctuations..

**Table 2.** Association between individual-specific S-A alignment and differences in age, sex, and puberty.

|  | Cortical Thickness |  | Functional Fluctuations |  | Isotropic Diffusion |  | Directional Diffusion |  |
| --- | --- | --- | --- | --- | --- | --- | --- | --- |
|  | Estimate | 95% CI | Estimate | 95% CI | Estimate | 95% CI | Estimate | 95% CI |
| Conditional $R^2$ | 0.119 | | 0.088 | | 0.196 | | 0.049 | |
| Marginal $R^2$ | 0.015 | | 0.009 | | 0.023 | | 0.008 | |
| Model $N$ | 2950 | | 2165 | | 2557 | | 2620 | |
| Age | -0.0343 | [-0.0733 – 0.0048] | -0.0178 | [-0.0643 – 0.0288] | **0.0543** | **[0.0123 – 0.0962]** | 0.0276 | [-0.0145 – 0.0697] |
| Age<sup>2</sup> | 0.0189 | [-0.0220 – 0.0597] | 0.0007 | [-0.0484 – 0.0498] | **0.0530** | **[0.0089 – 0.0970]** | -0.0059 | [-0.0499 – 0.0380] |
| Sex [Male] | **0.1384** | **[0.0394 – 0.2374]** | 0.0358 | [-0.0807 – 0.1522] | -0.0037 | [-0.1090 – 0.1015] | -0.0206 | [-0.1259 – 0.0847] |
| Puberty stage | **0.1425** | **[0.0779 – 0.2072]** | 0.0138 | [-0.0601 – 0.0878] | **0.1346** | **[0.0663 – 0.2028]** | 0.0105 | [-0.0579 – 0.0790] |
| Pubertal tempo | **0.1505** | **[0.0842 – 0.2168]** | -0.0120 | [-0.0880 – 0.0640] | 0.0679 | [-0.0022 – 0.1381] | 0.0058 | [-0.0647 – 0.0762] |
| Puberty stage x Sex [Male] | -0.0793 | [-0.1790 – 0.0204] | -0.0143 | [-0.1332 – 0.1047] | -0.0712 | [-0.1779 – 0.0355] | 0.0572 | [-0.0499 – 0.1642] |
| Pubertal tempo x Sex [Male] | **-0.1652** | **[-0.2487 – -0.0817]** | -0.0264 | [-0.1239 – 0.0710] | -0.0227 | [-0.1117 – 0.0663] | 0.0351 | [-0.0542 – 0.1244] |

*Note.* These report regression coefficients and associated 95% confidence intervals for linear mixed effects models of the form shown in Equation 3. Bold values indicate significant predictors at $p_{FWE} < 0.05$ ; bold and italicized values indicate significant predictors at $p_{FWE} < 0.01$ . For results of APA S-A alignment regressed on age, sex, puberty, and puberty by sex interactions, see Supplementary Table 7.

#### Neuroanatomical patterning of brain changes

Building off our previous work examining developmental changes in each region of the cortex, we used two metrics to estimate neurodevelopment over time, including: annualized percent change (APΔ) (Mills et al., 2021; Bottenhorn et al., 2023) and the reliable change index (RCI) over time (RCT) which assumes different variances at each time point (Maassen, 2004). In humans, APΔ has been previously used to describe structural and functional development and aging (Fjell et al., 2015; Walhovd et al., 2015; Tamnes et al., 2017; Mills et al., 2021; Bottenhorn et al., 2023), while RCI is more often used to assess clinical treatment effects (Duff, 2012; Blampied, 2022). Here, we extend the reliable change index to account for differences in inter-scan interval in scaling by the time elapsed between imaging time points to, instead, estimate reliable change over time. While APΔ accounts for differences in size or quality of the measurement (i.e., dividing by the average value between measurements), reliable change accounts for error in each measurement (i.e., by dividing by the standard errors, assuming independent variance per measurement; (Jacobson and Truax, 1992; Maassen, 2004)).

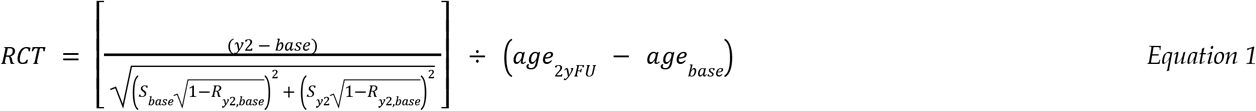

#### Age-related and individual differences in neurodevelopmental patterns

To assess participant alignment with the S-A axis across cortical thickness microarchitecture, and resting–state functional brain changes, we investigated both group-level and individual differences.

To probe group-level S-A associations with age, we performed ordinary least squares regressions using pinguoin (v0.5.1) in Python (Vallat, 2018). This aligns with prior work assessing differences in age slopes across the S-A axis (Sydnor et al., 2023). In doing so, we calculated Spearman correlations between individual-level brain changes (i.e., APΔ or RCT) and age for each brain region, across each aspect of neurodevelopment, then regressed those correlations on both linear and quadratic terms for regional S-A ranks (Equation 2).

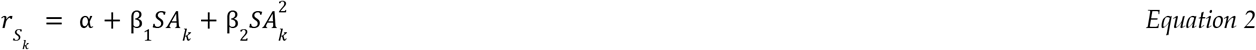

In Equation 2, *SA*_*k*_ is the S-A axis loading for the *k*^th^ brain region and *r*_*S,k*_ is the Spearman correlation between brain changes and age.

To assess the alignment of brain changes along the S-A axis, we computed Spearman correlations between each individual’s brain changes (i.e., APΔ or RCT) across regions with regions’ ranking along the S-A axis for cortical thickness, intracellular diffusion, and functional fluctuations Thus, an estimate of “S-A axis alignment” was calculated (i.e., as a Spearman correlation) for each individual, for each of the four MRI phenotypes assessed here (i.e., cortical thickness, isotropic intracellular diffusion, directional intracellular diffusion, and functional fluctuations). Due to the large variability of participant alignment with the S-A axis, next, we looked for differences in within-individual S-A alignment across sections of the S-A axis, to assess whether expected S-A patterning may still exist for portions of the axis. To do so, we calculated Spearman correlations between individual-level brain change (i.e., APΔ or RCT) and S-A axis rank per quartile along the S-A axis (e.g., within the regions comprising the most sensorimotor quarter of the S-A axis, etc.). We then compared individual-level alignment across sections of the S-A axis, within each aspect of neurodevelopment.

Lastly, we investigated if several developmental factors account for variation of individual differences seen inS-A alignment. To do so, we performed linear mixed effects models using the lme4 package (v1.1-29) (Bates et al., 2024) in R (v4.2.0), nesting participants within families and sites, while accounting for sociodemographic factors (Eq. 2). In these models, we were specifically interested in effects of age, sex, and pubertal timing and tempo, as well as if the latter two differed by sex.

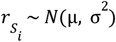, where *r*_*S, i*_ indicates S-A alignment of individual *i*’s brain changes, as a Spearman correlation.

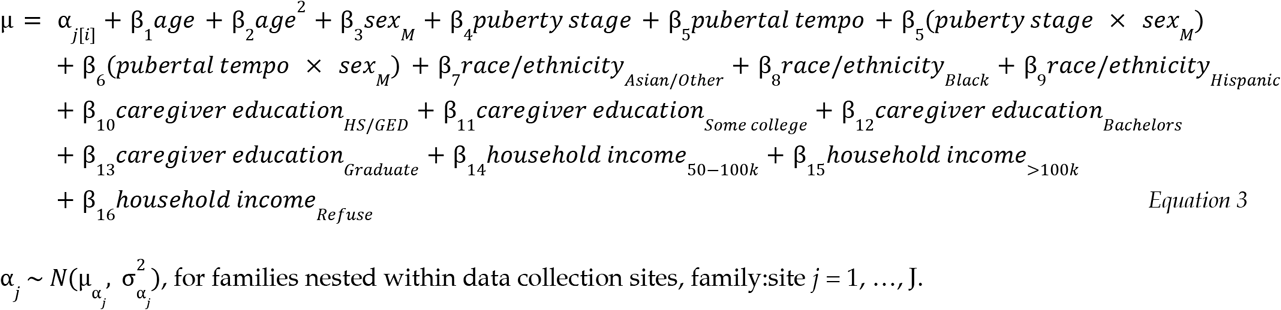

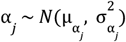, for families nested within data collection sites, family:site *j* = 1, …, J.

Simple slopes of any significant interaction were probed using the interaction package (v1.1.4) in R (Bauer and Curran, 2005; Long, 2021).

## Results

Across analyses, both metrics of brain change revealed similar trends, but reliable change over time (reliable change) was more sensitive to S-A organization than was annualized percent change (APΔ). This was evidenced by a greater proportion of individuals exhibiting statistically significant alignments of reliable change and the S-A axis as compared to APΔ. Thus, the results reported here focus on reliable change (hereafter: “change”), while all analyses performed with APΔ can be found in the Supplementary Materials.

### Group-level age effects along the S-A axis across measures of brain development

Across aspects of neurodevelopment, regional Spearman correlations between neurodevelopmental change and participants’ ages at 9 and 10 years exhibit both linear and quadratic progressions along the S-A axis (**Figure 1**, Supplementary Table 3). For cortical thickness, changes with age were largely negative: older individuals (i.e. 10 year-olds) demonstrated greater cortical thinning over time across most brain regions than did younger individuals (i.e., 9 year-olds). In observing how these changes with age vary by brain regions along the S-A axis, both linear and quadratic terms were significant suggesting a quadratic effect of age with an underlying S-to-A decrease. At both poles of the S-A axis (i.e., in the most sensorimotor- and association-biased regions) changes with age were largely zero, or even positive, while regions integrating sensorimotor and associative processing exhibited more marked decreases in cortical thickness with age. Changes in resting-state functional fluctuations were positively related to participants’ age, overall, and changes with age were found to follow a quadratic, inverted U-shaped pattern along the S-A axis. Changes in functional fluctuations with age were largely positive in most sensorimotor regions, followed by an increase in functional change with age towards the middle of the S-A axis, and with functional changes with age close to zero in more associative brain regions. Isotropic, but not directional, intracellular diffusion showed positive, linear associations between age-related brain changes and S-A axis loadings, such that older participants had greater decreases in isotropic intracellular diffusion within sensorimotor regions, but greater increases with age in association regions.

### Individual differences in S-A alignment across measures of brain development

Individual-level alignment of changes in cortical thickness, resting-state functional fluctuations, isotropic intracellular diffusion, and directional intracellular diffusion, were calculated by correlating regional reliable change scores with S-A axis loadings for each individual. Results showed divergent patterns of change across the cortex. Specifically, the magnitude of brain changes along the S-A axis ranged from *r*_*S*_ = 0.78 to *r*_*S*_ = -0.81 (Supplementary Figure 2A; Supplementary Tables 4), with the majority of individual’s showing an absence of S-A alignment (i.e., *r*_*S*_ =0). Across the sample, cortical thickness changes had an average S-A axis alignment of *r*_*S*_ *=* -0.004 (standard deviation, SD: 0.242; range: 1.551); functional fluctuations, an average of 0.037 (SD: 0.292; range: 1.594); isotropic diffusion, an average of -0.003 (SD: 0.251; range: 1.452); and directional diffusion, an average of 0.004 (SD 0.250; range: 1.487). Furthermore, an individual’s alignment with the S-A axis on one measure of brain change was unrelated to their alignment with the S-A axis on others (Supplementary Figure 2B). The same patterns were observed when individual-level alignment was assessed within smaller sections (i.e. each quartile) of the S-A axis (data not shown). For each measure of brain change, there were similar numbers of individuals with significant positive and negative correlations (**Figure 2**; Supplemental Figure 3; Supplementary Table 4). Using cortical thickness as an example, 11% of the sample showed greater cortical thinning in association regions than in sensorimotor regions, while 10% showed greater cortical thinning in sensorimotor regions than in association regions (**Figure 3C**). Similar patterns were observed in isotropic intracellular diffusion changes (Figure 3D), as well as changes in functional fluctuations and directional intracellular diffusion (Supplemental Figure 3; Supplementary Table 4).

**Figure 2.**
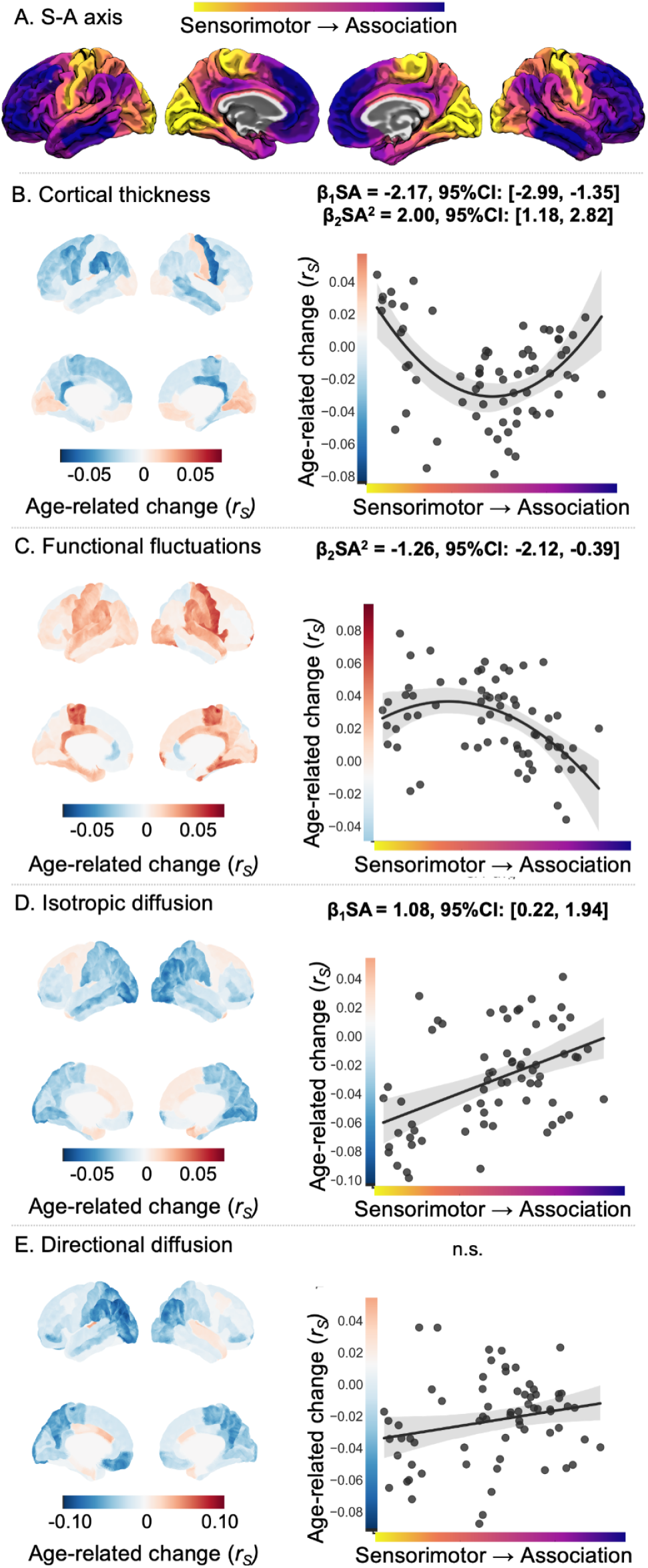
Correlations between reliable change and participant age are different across the S-A axis. A. Reference schematic of the S-A axis, created with data from Sydnor et al. (2021). In B-E, brain images (right) show region-wise change in brain development with age, whereas graphs (left) depict brain changes with age (y-axis)) regressed along S-A axis loadings (x-axis) for various neurodevelopmental outcomes, including B.) cortical thickness; C.) resting-state functional fluctuations; D.) isotropic intracellular diffusion; and E.) directional intracellular diffusion. For graphs in B-E, each point represents one cortical region, x-axis is color-coded according to S-A rank (see: A) and y-axis is color-coded according to age-change correlations (see: corresponding brain images). Bold standardized coefficients (i.e., *β*_1_*SA* and *β*_2_*SA*^2^; see Equation 2) indicate significant associations (*a*_*FWE*_ < 0.015) in age-related change in brain development along the S-A axis. All coefficients, confidence intervals, and associated *p*-values are in Supplementary Table 3.

**Figure 2.**
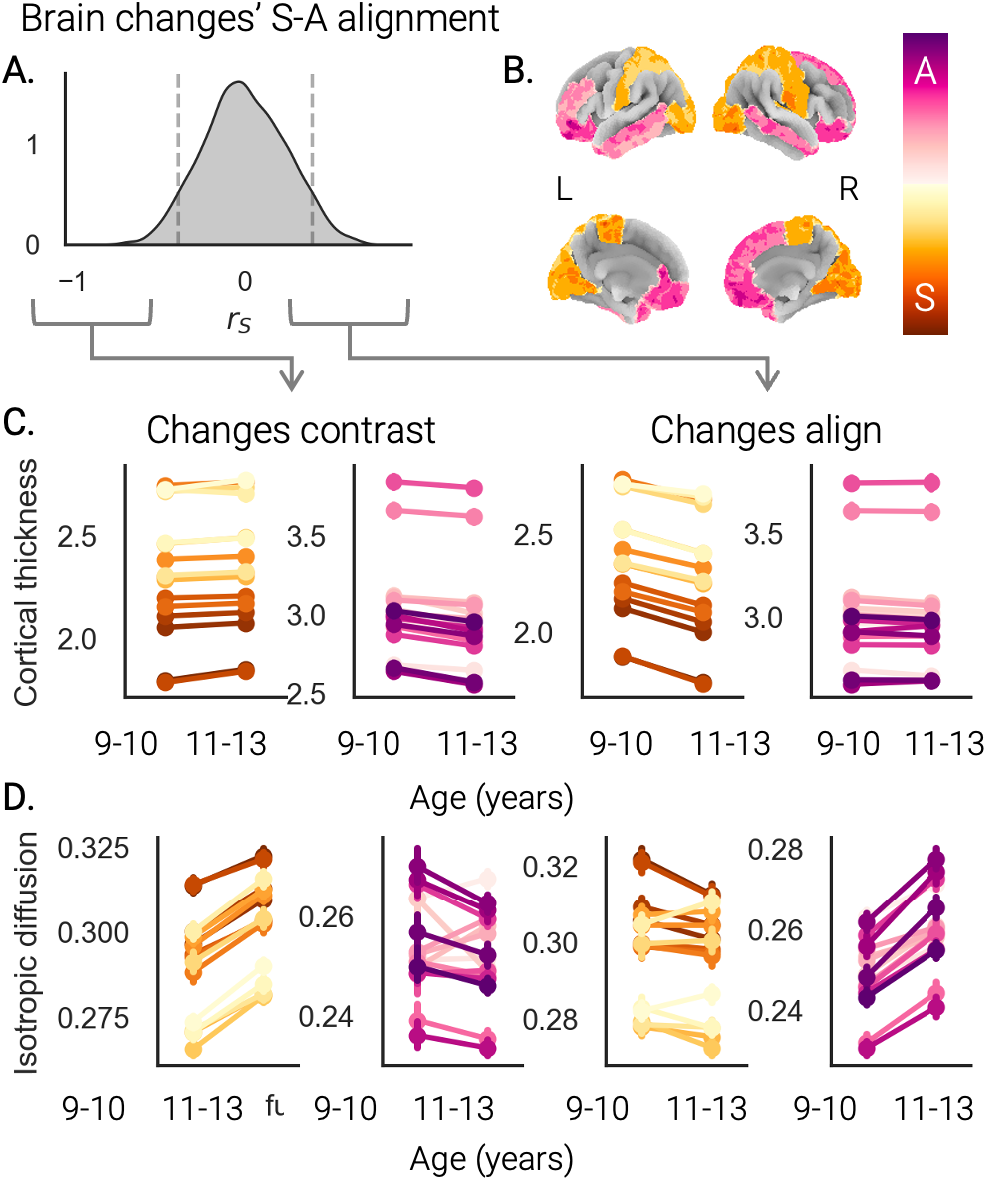
Individual-level alignment of cortical thickness and isotropic diffusion changes along the S-A axis. Spearman correlations (*r*_*S*_) between brain changes in each region and S-A axis loadings revealed widespread individual differences in spatiotemporal patterns of cortical development between 9 and 13 years of age. (A) Here, we present the distribution of how individual-level changes in cortical thickness align with the S-A axis with dashed lines denoting the cutoffs for “significant” correlations. To illustrate the variety in S-A patterning of brain changes (i.e., *r*_*S*_), we plot the brain change scores for (B) the most sensorimotor (yellow-orange) and most associative (pink-purple) brain regions for (C) cortical thickness and (D) isotropic intracellular diffusion. (C) Individuals with significant negative S-A alignment in cortical thickness displayed less cortical thinning in sensorimotor as compared to associative brain regions (left), whereas individuals with significant positive S-A alignment in cortical thickness displayed greater cortical thinning in sensorimotor regions from ages 9-10 years as compared to association areas that showed little to no change over this 2-year follow-up period (left) (D) Individuals with significant negative S-A alignment in isotropic intracellular diffusion showed greater increases in sensorimotor regions as compared to no changes and decreases in associative brain regions (left), whereas those displaying a significant positive alignment showed the opposite pattern of changes over the 2-year period (i.e. changes were zero or negative for sensorimotor regions; increases were seen for associative brain regions S-A alignment values for cortical thickness. Error bars in C and D indicate the 95% confidence interval about the mean. Corresponding graphs for functional fluctuations and directional diffusion can be found in Supplementary Figure 3.

### Individual differences in S-A alignment relate to age, sex, and pubertal maturation

To better understand variability in participant alignment with the S-A axis, we examined if individual’s alignment scores (i.e., participant-wise correlation with S-A axis; *r*_*S*_) were associated with age, sex, pubertal timing, pubertal tempo, as well as possible interactions between puberty and sex (i.e., pubertal timing-by-sex; pubertal tempo-by-sex; see Methods Eq. 2). After correcting for the number of effective comparisons between brain measures (α < 0.05, *M*_*eff*_ = 3.33: *p*_*adj*_ = 0.0152), we found that age, sex, and pubertal development explain some of the variation in individual-level S-A alignment (**Table 2**). For cortical thickness, individual-level S-A alignment scores were found to be significantly associated with pubertal timing and tempo as a function of sex (i.e. pubertal timing-by-sex and pubertal tempo-by-sex interactions; **Figure 4**). Comparing the slopes of this interaction suggests a significant, positive association between S-A alignment and pubertal tempo for female youth (B = 0.10, SE = 0.02, 95% CI = [0.06, 0.14], *p* < 0.001), but no significant association for male youth (B = -0.01 SE = 0.02, 95% CI = [-0.04, 0.02], *p* = 0.60; Figure 4E). For functional fluctuation changes (Figure 4B), individual differences in S-A alignment was not related to age, sex, or pubertal development. For isotropic diffusion changes (Figure 4C), individual differences in S-A alignment showed a significant quadratic effect with age, were greater in male youth and youth with both later pubertal timing and quicker pubertal tempo. For directional diffusion (Figure 4D), individual differences in S-A alignment were unrelated to age, sex, or pubertal development.

**Figure 4.**
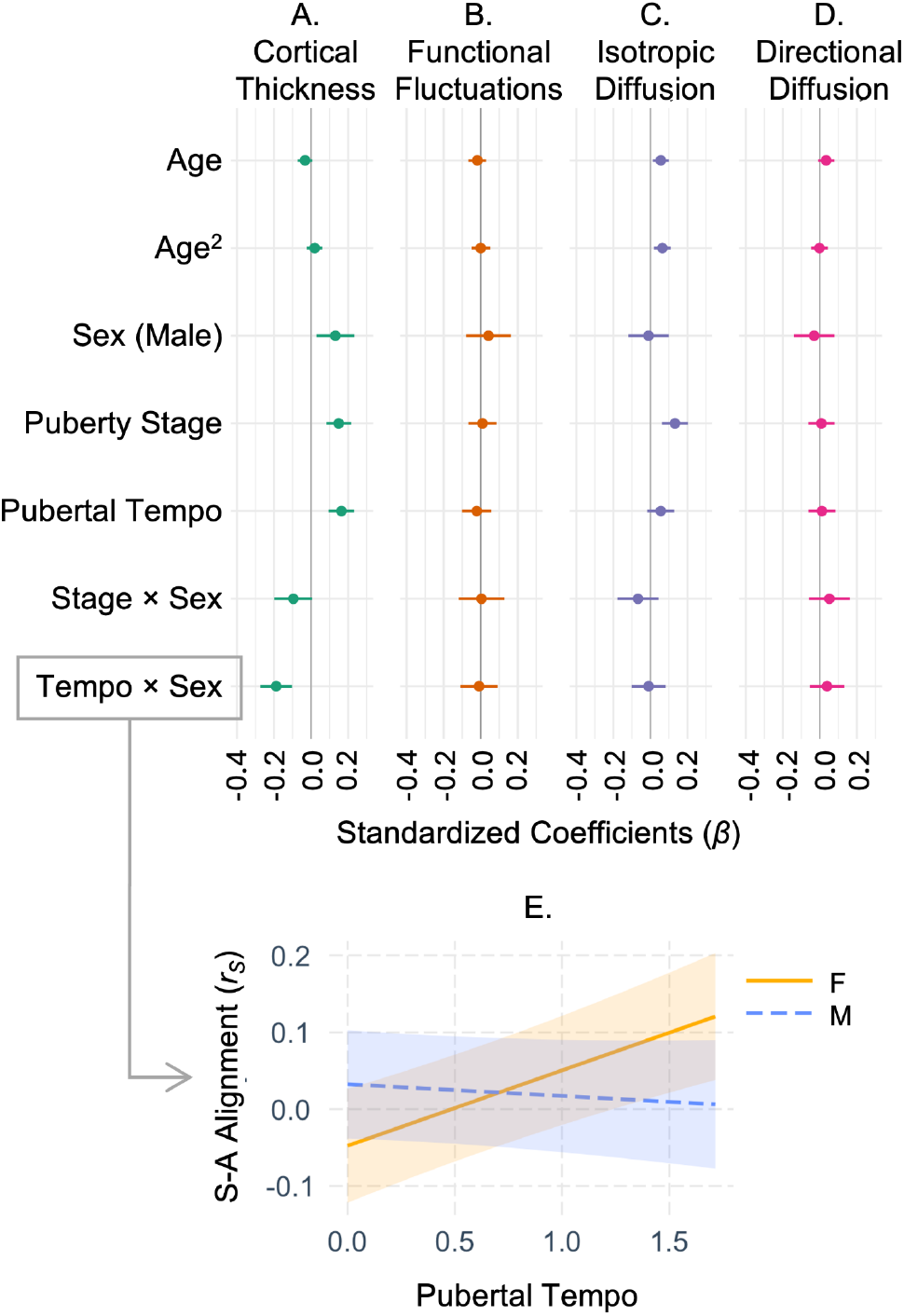
Relationships between individual differences in S-A alignment and age, sex, and puberty. (A) Individual differences in alignment of cortical thickness changes along the S-A axis are associated with pubertal timing and tempo as a function of sex (i.e. interactions). (B) Individual differences changes in resting-state functional fluctuations along the S-A axis are not related toage, sex, or puberty. (C) Individual differences in isotropic intracellular diffusion changes along the S-A axis are associated with age, pubertal tempo, and pubertal timing as a function of sex (i.e., interaction). (D) Individual differences in directional intracellular diffusion changes along the S-A axis are not related to age, sex, or puberty. (E) Simple slopes for interactions of sex (female, F; male, M) and pubertal tempo (x-axis) related to individual-level S-A alignment of cortical thickness changes (*r*_*S* ;_ y-axis). For numerical values, see Table 2.

## Discussion

This work investigated patterns of early adolescent changes in cortical thickness, function (i.e., BOLD fluctuations), and microarchitecture (i.e., isotropic, directional intracellular diffusion). Brain regions’ positioning along a sensorimotor-association (S-A) axis partially explained group-level, age-related changes in cortical thickness, functional fluctuations, and isotropic intracellular diffusion (estimating cellular density). However, individual differences in brain changes along the S-A axis were considerable; most youth did not exhibit S-A organization in cortical changes. Depending on the MRI phenotype, only 21-33% of early adolescents’ brain changes progressed along the S-A axis: approximately half of these individuals’ regional brain changes increased from sensorimotor to association areas, while the other half decreased. Individual-specific S-A alignment of cortical thickness and cellular density changes varied with age, sex, and puberty. Altogether, S-A patterning in early adolescent cortical development seems to include both age-related, between-individual differences and marked within-individual variability.

### Age differences in developmental change

Human developmental neuroimaging research often relies on cross-sectional data, conflating age-related variance and true developmental change. Cross-sectional data can yield sound longitudinal inferences only if trajectories are linear and measurement error is constant with age (Kraemer et al., 2000). However, brain development exhibits curvilinear age-related changes (Wierenga et al., 2014; Bethlehem et al., 2022) and the degree of motion-related noise in MRI data varies with age (Satterthwaite et al., 2012; Madan, 2018). Recent analyses of ABCD Study neuroimaging data show cross-sectional models poorly predict individual developmental change, as individual differences in development can outweigh age-related variance (Di Biase et al., 2023). Thus, this work lends longitudinal insight to a largely cross-sectional literature by assessing S-to-A patterning of *changes* in several MRI phenotypes, using two imaging acquisitions, at ages 9-10 years and 11-13 years, from thousands of individuals. In cortical thickness, prior human MRI studies suggest earlier gray matter maturation in sensorimotor regions than in associative regions (Gogtay et al., 2004; Kalantar-Hormozi et al., 2023). The present study identified more nuanced patterns in early adolescence; age-related cortical thinning was greatest in the middle of the S-A axis. This may indicate that, from ages 9-13 years, cortical maturation is in the midst of a larger S-to-A progression: cortical thinning has slowed in primary sensorimotor regions but barely begun in association regions. Supporting this notion are findings that early adolescent cortical thinning is least in primary sensory and prefrontal cortices (i.e., the poles of the S-A axis), but greatest in precuneus, paracentral gyrus, and superior parietal lobule, (i.e., the middle of the S-A axis) (Wierenga et al., 2014; Tamnes et al., 2017). Intrinsic brain activity has shown negative age differences in sensorimotor regions, but positive age differences in association regions at 10 years of age (Sydnor et al., 2023); functional connectivity increases from age 7 to 13 years in transmodal networks, but decreases in sensory networks (Xia et al., 2022). Here, we identified a curvilinear pattern; functional fluctuations increase with age most in predominantly sensorimotor regions, but decrease or remain stable in more associative brain regions. Finally, greater age-related increases in cortical myelination have been identified in sensorimotor regions than association regions (Baum et al., 2022). Here, we used novel metrics from isotropic and directional intracellular diffusion to estimate cellular density (e.g., neuronal and support cell bodies; reflecting apoptosis, neurogenesis, inflammation) and neurite density (e.g., axons and dendrites; reflecting myelination, synaptic refinement), respectively. We observed sensorimotor decreases and associative increases in age-related cellular density changes, but no significant S-A patterning in age-related neurite density changes.

### Individual variability in spatiotemporal patterns of cortical development

Normative models can provide foundations for understanding healthy and abnormal neurodevelopment (Bethlehem et al., 2022; Rutherford et al., 2023), but can overlook considerable, heterogeneous individual differences (Mills et al., 2021; Bottenhorn et al., 2023). Neural variability may be greater in early development (Gratton et al., 2018; Wang et al., 2021) and in association cortex (Mueller et al., 2013; Laumann et al., 2015; Seitzman et al., 2019; Cui et al., 2020), potentially compromising the accuracy of normative models and of our understanding of S-to-A neurodevelopmental patterning. Our individual-centric approach addresses such drawbacks.

Here, we show that S-A patterns of early adolescent cortical development vary considerably between individuals and within individuals, across aspects of development. Individuals exhibiting greatest cortical thinning in sensorimotor regions and least, in association regions (i.e., positive S-A alignment) do not necessarily display S-A alignment in cortical microarchitecture or functional changes. This disconnect may reflect timing and tempo differences in neurobiological mechanisms underlying different aspects of development. Further, this narrow age range (9-13 years) likely presents an incomplete picture of S-A neurodevelopmental patterns, which, according to prior, cross-sectional work in a wider-age range, peaks at 14.5 years of age (Sydnor et al., 2023).

Puberty’s role in S-to-A neurodevelopmental patterning is unclear. Grydeland et al. found that puberty bisects cortical myelination peaks in sensorimotor regions and in association, insular, and limbic regions (Grydeland et al., 2019), but Sydnor et al. found no significant association between puberty and S-A patterns in intrinsic activity, beyond age differences (Sydnor et al., 2023). Our individual-centric approach uncovered sex-moderated roles of pubertal development in early adolescent cortical thinning patterns along the SA axis. Greater S-A alignment of cortical thinning was observed in youth who were farther along in puberty at ages 9-10 years, which was stronger for female than male youth such that, with faster pubertal tempo, female youth showed greater positive S-A alignment; male youth, greater negative alignment. As pubertal development occurs earlier in female than male youth, we may observe similar pubertal influences on S-A patterning in male youth at later ages. In cellular density changes, youth who were older, male, and who exhibited earlier puberty and faster pubertal tempo, regardless of sex, showed greater positive S-A alignment. If deviations from an S-to-A pattern of cortical development are, indeed, related to psychopathology risk (Gogtay et al., 2004; Sydnor et al., 2021), then further investigating these findings may explicate neuroendocrine sex differences in adolescent psychopathology (Martel, 2013).

### Limitations

This insight into spatiotemporal organization of early adolescent cortical development in a large sample faces some limitations. Primarily, the ABCD Study’s biennial imaging assessments may insufficiently capture the spatiotemporal organization of cortical change during this dynamic developmental period. Second, both metrics used here to estimate neurodevelopment (i.e., APΔ and RCT) have drawbacks. APΔ with two time points cannot account for measurement error, while the RCI denominator conflates individual variability and measurement error, potentially underestimating individual differences. Results converged between metrics, although RCT, but not APΔ, revealed significant S-A patterns of age-related change, greater individual variability, and greater sensitivity to differing age, sex, and pubertal development. Generally, two imaging assessments per individual can model individual-level change, but cannot capture nonlinearities in change or within-individual error. While adding additional time points to this work would better characterize individual-level change, the global COVID-19 pandemic occurred between most ABCD Study youth’s 2-year and 4-year imaging assessments (i.e., between ages 11-13 years and 13-14 years), introducing considerable, unknown effects on adolescent development. Future research with the ABCD Study should develop strategies for mitigating and understanding confounding pandemic experiences on adolescent development. Broadly, precision neuroscience research should investigate measurement error, and age-related changes therein, to disentangle error from meaningful individual differences, where possible.

This operationalization of perceived pubertal timing and tempo uses Tanner stage estimates based on caregiver-reported perceived physical changes on the Pubertal Development Scale (PDS; (Petersen et al., 1988)). Prior work shows that PDS scores correlate with sex steroid hormone changes and that dehydroepiandrosterone, estradiol, and testosterone levels significantly differ with PDS scores (Herting et al., 2021). However, PDS scores represent *perceived* physical changes accompanying pubertal development, providing less precise Tanner stage estimates than do physician reports. We estimated Tanner stages due to their prevalence in the pubertal development literature. Unfortunately, the PDS provides less information about the earlier stages of puberty than later (Cheng et al., 2021), potentially underscoring some sex-moderated pubertal differences in neurodevelopmental patterning identified here. Thus, while this work provides preliminary evidence of pubertal differences in early adolescent neurodevelopmental patterning, future work should incorporate complementary measures of pubertal development, such as sex steroid hormone levels and height velocity (Herting et al., 2021; Byrne et al., 2023).

Although the ABCD Study aimed to recruit a nationally-representative sample in the United States, the final baseline sample overrepresented educated and affluent families (Dick et al., 2021; Simmons et al., 2021), but underrepresented individuals of Asian, American Indian/Alaska Native and Native Hawaiian/Pacific Islander ancestry (Heeringa & Berglund, 2020). MRI quality control further narrowed the sample unequally across demographics, additionally limiting the generalizability.

## Conclusions

While group- and individual-level changes in cortical morphology, microarchitecture, and function between ages 9 and 13 years show some sensorimotor-to-association organization, individual-level S-A patterning in cortical development is remarkably heterogeneous. Differential patterns of individual-level change in early adolescence may highlight varying morphological, functional, and microarchitectural trajectories, with potential influences from age, sex, and pubertal development. Overall, this work highlights diverse neurodevelopmental trajectories underscored by age-related variance and individual-specific pubertal factors, with both within- and between-individual variation that requires further study.

## Acknowledgments

A special thank you to all the children and families for their participation in their ABCD Study.

Research described in this article was supported by the National Institutes of Health (MMH: NIEHS R01ES032295, R01ES031074; KLB: P30ES07048-27, K99MH135075).

Data used in the preparation of this article were obtained from the Adolescent Brain Cognitive Development^SM^ (ABCD) Study (https://abcdstudy.org), held in the NIMH Data Archive (NDA). This is a multisite, longitudinal study designed to recruit more than 10,000 children age 9-10 and follow them over 10 years into early adulthood. The ABCD Study® is supported by the National Institutes of Health and additional federal partners under award numbers U01DA041048, U01DA050989, U01DA051016, U01DA041022, U01DA051018, U01DA051037, U01DA050987, U01DA041174, U01DA041106, U01DA041117, U01DA041028, U01DA041134, U01DA050988, U01DA051039, U01DA041156, U01DA041025, U01DA041120, U01DA051038, U01DA041148, U01DA041093, U01DA041089, U24DA041123, U24DA041147. A full list of supporters is available at https://abcdstudy.org/federal-partners.html. A listing of participating sites and a complete listing of the study investigators can be found at https://abcdstudy.org/consortium_members/. ABCD consortium investigators designed and implemented the study and/or provided data but did not necessarily participate in the analysis or writing of this report. This manuscript reflects the views of the authors and may not reflect the opinions or views of the NIH or ABCD consortium investigators. The ABCD data repository grows and changes over time. The ABCD data used in this report came from http://dx.doi.org/10.15154/z563-zd24.

## Author Contributions

Conceptualization: KLB, MMH

Data curation: KLB

Formal Analysis: KLB, JDC

Funding acquisition: MMH

Methodology: KLB, MMH, HA

Project administration: KLB, MMH

Resources: MMH

Software: KLB

Supervision: MMH

Visualization: KLB

Writing – original draft: KLB, JDC, MMH

Writing – review & editing: KLB, JDC, MMH

