## Supplementary Material for "Spatiotemporal patterns in cortical development: Age, puberty, and individual variability from 9 to 13 years of age"

This document includes:

- Supplementary Methods
- Supplementary Results
- Supplementary Tables 1 to 7
- Supplementary Figures 1 to 3

#### Supplementary Methods

##### *Participants: QC & Demographics*

Data cleaning and quality control restricted the sample included in these analyses to individuals with sufficient quality MRI data collected at the baseline (i.e., ages 9-10 years) and 2-year follow-up (i.e., ages 10-13 years) visits, with 2-year follow-up visits prior to the onset of the COVID-19 pandemic (i.e., March 2020), and all MRI data collected on a SIEMENS 3T scanner. Below in Supplementary Table 1 are the demographics of the participants who were excluded from the analyses presented in this work, alongside the whole sample demographics for comparison.

##### *Missing Data*

Neuroimaging data was missing at random with respect to sociodemographic factors, such that youth from lower-income households had more missing data and non-white youth had more missing data. Female youth had less missing fMRI data, but more missing sMRI data compared to male youth in this sample. Finally, neuroimaging data missingness was not equally distributed across study sites, which was furthered by restricting our sample to data collected on SIEMENS MRI scanners. Amounts of missing data on neuroimaging, developmental, and demographic data are shown below in Supplementary Table 2.

*Corresponding Author*: Dr. Katherine L. Bottenhorn, Postdoctoral Scholar, Department of Population and Public Health Sciences, University of Southern California, Los Angeles, CA USA |

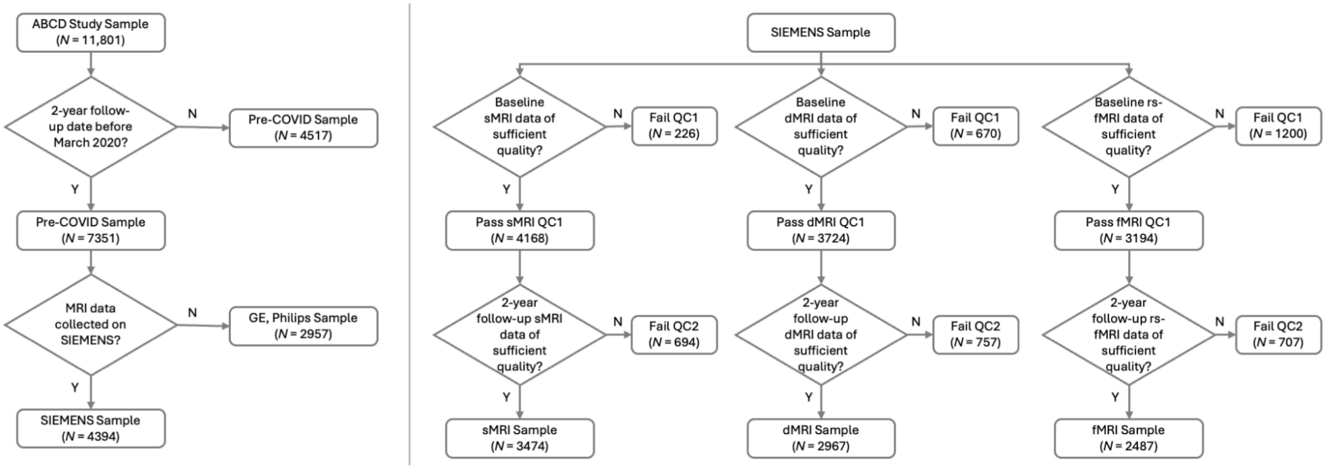


***Supplementary Figure 1. Participant inclusion flow chart.*** From the full ABCD Study sample (*N* = 11,868), this study included participants whose 2-year follow-up visit was before March 2020 (*N* = 7,351; excluded *N* = 4,517) with MRI data acquired on a SIEMENS scanner (*N* = 4,394; excluded *N* = 2,957). Of these participants, imaging data were included if both baseline and 2-year follow-up MRI scans were of sufficient quality (see *Methods* for details).

**Supplementary Table 1.** Total and excluded participant demographics.

| | Full ABCD Study Sample | | Post-COVID Follow-up | | Cortical Thickness Excluded | | Intracellular Diffusion Excluded | | Functional Fluctuations Excluded | |
|---|---|---|---|---|---|---|---|---|---|---|
| | N | % of sample | N | % of sample | N | % of sample | N | % of sample | N | % of sample |
| Total N | 11868 | 100% | 4517 | 38% | 8394 | 71% | 8901 | 75% | 9381 | 79% |
| Age at baseline (months) | 118.98 ± 7.496 | 1% | 118.25 ± 7.52 | 1% | 118.76 ± 7.53 | 1% | 118.77 ± 7.51 | 1% | 118.73 ± 7.47 | 1% |
| Sex (F) | 5677 | 48% | 2184 | 18% | 4081 | 34% | 4282 | 36% | 4436 | 37% |
| Puberty | | | | | | | | | | |
| Pre-pubertal | 5837 | 49% | 2117 | 18% | 4068 | 34% | 4297 | 36% | 4555 | 38% |
| Early puberty | 2709 | 23% | 1010 | 8% | 1895 | 16% | 2016 | 17% | 2128 | 18% |
| Mid-puberty | 2672 | 23% | 1057 | 9% | 1920 | 16% | 2047 | 17% | 2141 | 18% |
| Late/post puberty | 181 | 2% | 85 | 1% | 138 | 1% | 144 | 1% | 150 | 1% |
| Puberty change | | | | | | | | | | |
| Puberty | 0.41 ± 0.42 | 0% | 0.42 ± 0.39 | 0% | 0.41 ± 0.42 | 0% | 0.41 ± 0.42 | 0% | 0.41 ± 0.42 | 0% |
| Puberty Missing | 2403 | 20% | 1511 | 13% | 1974 | 17% | 2060 | 17% | 2131 | 18% |
| Race & Ethnicity | | | | | | | | | | |
| Asian + Other | 1499 | 13% | 614 | 5% | 1165 | 10% | 1228 | 10% | 1284 | 11% |
| Hispanic | 2410 | 20% | 996 | 8% | 1811 | 15% | 1896 | 16% | 1983 | 17% |
| Non-Hispanic Black | 1784 | 15% | 908 | 8% | 1319 | 11% | 1430 | 12% | 1526 | 13% |
| Non-Hispanic White | 6173 | 53% | 1997 | 17% | 4097 | 35% | 4345 | 37% | 4586 | 39% |
| Household Income | | | | | | | | | | |
| < $50,000 | 3222 | 27% | 1386 | 12% | 2389 | 20% | 2554 | 22% | 2695 | 23% |
| $50,000 to $100,000 | 3068 | 23% | 1065 | 9% | 2033 | 17% | 2179 | 18% | 2318 | 20% |
| > $100,000 | 4561 | 38% | 1599 | 13% | 3196 | 27% | 3347 | 28% | 3508 | 30% |
| Caregiver Education | | | | | | | | | | |
| Up to high school diploma, GED | 593 | 5% | 287 | 2% | 482 | 4% | 513 | 4% | 538 | 5% |
| High school diploma, GED | 1132 | 10% | 561 | 5% | 868 | 7% | 932 | 8% | 987 | 8% |
| Some college, associate's degree | 3074 | 26% | 1238 | 10% | 2150 | 18% | 2305 | 19% | 2443 | 21% |
| Bachelor's degree | 3013 | 25% | 1006 | 8% | 2014 | 17% | 2134 | 18% | 2268 | 19% |
| Graduate degree | 4042 | 34% | 1421 | 12% | 2872 | 24% | 3009 | 25% | 3134 | 26% |

**Supplementary Table 1.** Total and excluded participant demographics.

| | Full ABCD Study Sample | | Post-COVID Follow-up | | Cortical Thickness Excluded | | Intracellular Diffusion Excluded | | Functional Fluctuations Excluded | |
|---|---|---|---|---|---|---|---|---|---|---|
| | N | % of sample | N | % of sample | N | % of sample | N | % of sample | N | % of sample |

*Note*. Within-category percentages that add to <100% are due to missing data and participants who declined to answer, e.g., in the case of household income. Abbreviations: GED, General Educational Development test.

**Supplementary Table 2.** Missing data across variables.

| | Full ABCD Study Sample | | Cortical Thickness | | Cortical Intracellular Diffusion | | Resting-state Functional Fluctuations | |
|---|---|---|---|---|---|---|---|---|
| | N | % of sample | N | % of sample | N | % of sample | N | % of sample |
| Total N | 11868 | 100% | 3474 | 29% | 2967 | 25% | 2487 | 21% |
| Age | 0 | | 0 | | 0 | | 0 | |
| Sex | 0 | 0% | 0 | 0% | 0 | 0% | 0 | 0% |
| Puberty at baseline | 469 | 4% | 96 | 1% | 72 | 1% | 62 | 1% |
| Puberty at 1-year | 1199 | 10% | 243 | 2% | 194 | 2% | 146 | 1% |
| Puberty at 2-year | 1582 | 13% | 176 | 1% | 142 | 1% | 122 | 1% |
| Race & Ethnicity | 2 | 0% | 2 | 0% | 2 | 0% | 2 | 0% |
| Household Income | 2 | 0% | 2 | 0% | 2 | 0% | 2 | 0% |
| Caregiver Education | 14 | 0% | 8 | 0% | 8 | 0% | 11 | 0% |

### Supplementary Results

#### *Reliable change over time (RCT)*

**Supplementary Table 3.** Associations of age and brain changes regressed along the S-A axis.

| | Standardized Coefficient ($\beta$) | $p$ | 95% CI | $R^2_{Adj}$ |
|---|---|---|---|---|
| Cortical Thickness | | | | 0.287 |
| Linear term | -2.17 | **1.54 ×10$^{-6}$** | [-2.99, -1.35] | |
| Quadratic term | 1.99 | **7.62 ×10$^{-6}$** | [1.18, 2.82] | |
| Functional fluctuations | | | | 0.208 |
| Linear term | 0.85 | 0.0529 | [-0.01, 1.72] | |
| Quadratic term | -1.26 | **0.00505** | [-2.12, -0.39] | |
| Isotropic Diffusion | | | | 0.2176 |
| Linear term | 1.08 | **0.0143** | [0.22, 1.94] | |
| Quadratic term | -0.00003 | 0.142 | [-1.50, 0.22] | |
| Directional Diffusion | | | | 0.056 |
| Linear term | 0.95 | 0.048 | [0.01, 1.89] | |
| Quadratic term | -0.75 | 0.115 | [-1.70, 0.19] | |

***Note.*** Results reflect the linear and quadratic terms of the model. The associations noted represent the S-A axis loadings, onto which age associations with RC/T-age (i.e., correlations of age and RC/T per region, across participants) were regressed. No correlations between APΔ and age were significantly related to S-A axis loadings, either linearly or quadratically (Supplementary Table 5). Bold $p$-values are significant at $\alpha < 0.05$, adjusted for the number of effective comparisons between models (i.e., $p < 0.015$).

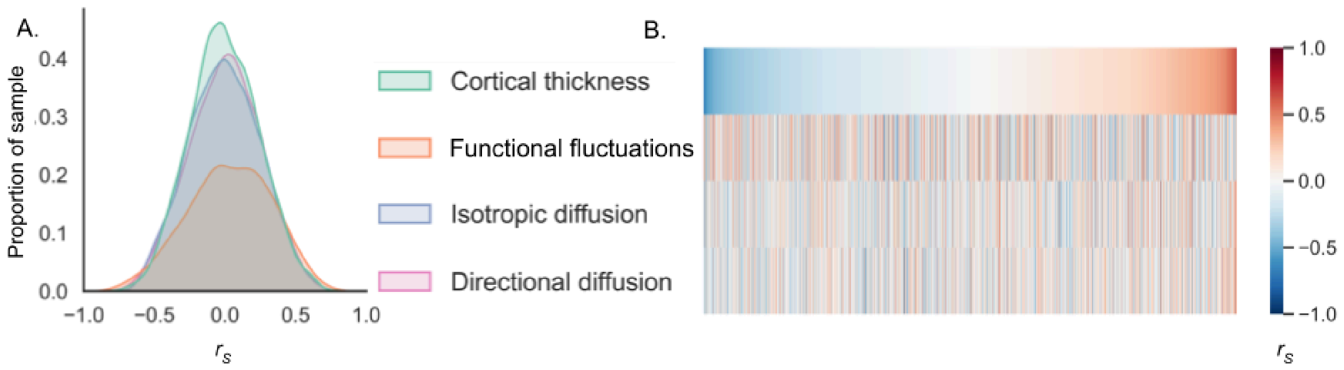


***Supplementary Figure 2. Individual-level alignment of brain changes along the S-A axis.*** Individual-level alignment with the S-A axis was computed via Spearman correlations of change in each brain region with regions' S-A axis ranks, within each individual. This provides an estimate of how each individual's pattern of change across the brain progresses from sensorimotor to associative regions, such that an individual showing greater increases in associative regions than in sensorimotor regions would demonstrate a positive alignment and an individual showing greater increases in sensorimotor regions than associative regions would demonstrate a negative alignment. A.) Distributions of participant alignments with the S-A axis, across change in each brain measure over time. B.) Each participant's Spearman correlation between regional change over time and S-A axis loadings is color-coded and represented in a column. Columns (participants) are ordered by their correlation between change in cortical thickness and the S-A axis (i.e., S-A axis alignment for cortical thickness).

**Supplementary Table 4.** Alignment of individual reliable change over time (RCT) with sensorimotor-association axis**.**

| | Average correlation | Standard deviation | Positive correlation *N* (%) | Negative correlation (N) |
|---|---|---|---|---|
| Cortical Thickness | -0.004 | 0.242 | 362 (10%) | 373 (11%) |
| BOLD Variance | 0.038 | 0.292 | 482 (19%) | 337 (14%) |
| Isotropic Diffusion | -0.003 | 0.251 | 336 (11%) | 336 (11%) |
| Directional Diffusion | 0.004 | 0.250 | 328 (11%) | 350 (12%) |

***Note.*** "Contrasted" refers to significantly negative Spearman correlations ($p <$ 0.01) between the individual's RCTs and their expected loading on the S-A axis. Corresponding results for APΔ are available in Supplementary Table 6.

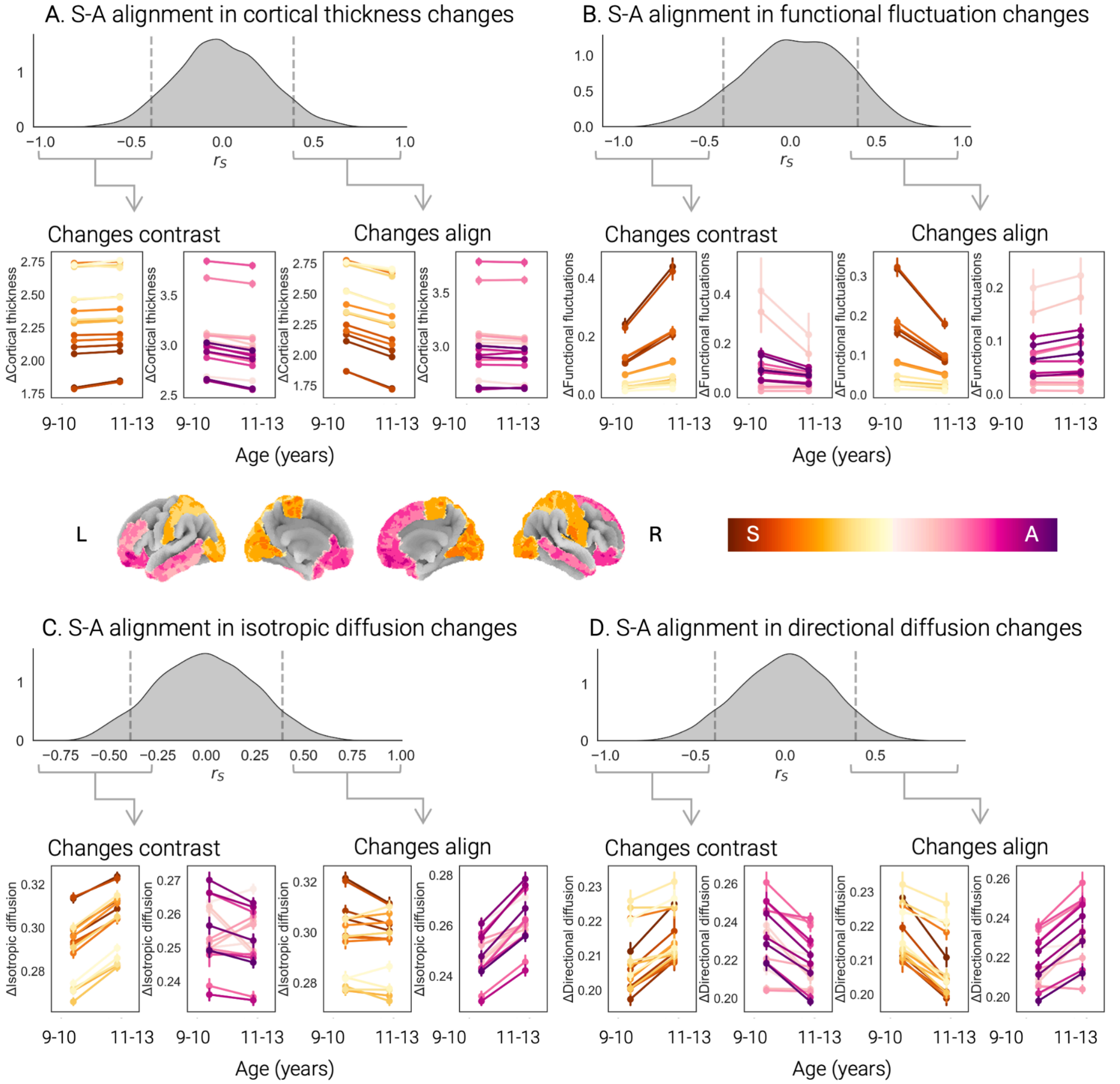


***Supplementary Figure 3. Variable individual-level alignment of RCT along the S-A axis.*** Spearman correlations ($r_S$) between regional brain changes and regional loadings along the S-A axis revealed widespread individual differences in spatiotemporal patterns of cortical development between 9 and 13 years of age. Here, we present the distribution of individual-level S-A alignment in cortical thickness changes (A), functional fluctuation changes (B), isotropic intracellular diffusion changes (C), and directional diffusion changes (D) with dashed lines denoting the cutoffs for "significant" correlations. S Here, to illustrate the variety in S-A patterning of brain changes in this sample, we show, for the most sensorimotor and most associative regions (visualized on the cortical surface in the center of the figure), average values of cortical thickness (A), functional fluctuations (B), isotropic intracellular diffusion (C) , and directional intracellular diffusion (D) at each imaging data collection time point (i.e., at ages 9-10 years and 11-13 years) across individuals with significantly negative S-A alignment and those with significantly positive S-A alignment. Error bars in point plots indicate the 95% confidence interval about the mean (plotted).

#### *Annualized percent change (APΔ)*

**Supplementary Table 5.** Associations of age and APΔ regressed along the S-A axis.

| | Standardized Coefficient ($\beta$) | $p$ | 95% CI | $R^2_{Adj}$ |
|---|---|---|---|---|
| Cortical Thickness | | | | 0.056 |
| Linear term | 0.73 | 0.128 | [-0.22, 1.67] | |
| Quadratic term | -0.48 | 0.314 | [-1.42, 0.46] | |
| Functional fluctuations | | | | 0.015 |
| Linear term | -0.27 | 0.575 | [-1.24, 0.69] | |
| Quadratic term | 0.46 | 0.341 | [-0.50, 1.43] | |
| Isotropic Diffusion | | | | -0.027 |
| Linear term | -0.10 | 0.832 | [-1.09, 0.88] | |
| Quadratic term | 0.05 | 0.925 | [-0.94 , 1.03] | |
| Directional Diffusion | | | | -0.013 |
| Linear term | -0.21 | 0.674 | [-1.18, 0.77] | |
| Quadratic term | 0.08 | 0.868 | [-0.90, 1.06] | |

***Note.*** Results reflect the linear and quadratic terms of the model. The associations noted represent the S-A axis loadings, onto which age associations with APΔ-age (i.e., correlations of age and APΔ per region, across participants) were regressed. Bold $p$-values are significant at $\alpha < 0.05$, adjusted for the number of effective comparisons between models (i.e., $p < 0.015$).

**Supplementary Table 6.** Alignment of individual annual percent brain changes with sensorimotor-association axis.

| | Average correlation | Standard deviation | Positive correlation | Negative correlation |
|---|---|---|---|---|
| Cortical Thickness | -0.006 | 0.160 | 88 (3%) | 101 (3%) |
| BOLD Variance | -0.064 | 0.191 | 55 (2%) | 237 (10%) |
| Isotropic Diffusion | 0.006 | 0.183 | 148 (5%) | 115 (4%) |
| Directional Diffusion | -0.004 | 0.159 | 70 (2%) | 86 (3%) |

***Note.*** "Contrasted" refers to significantly negative Spearman correlations ($p < 0.01$) between the individual's brain changes and their expected loading on the S-A axis, based on expected developmental trajectories.

**Supplementary Table 7.** Association between individual-specific S-A alignment (based on APΔ) and differences in age, sex, and puberty.

| | Cortical Thickness | | Functional Fluctuations | | Isotropic Diffusion | | Directional Diffusion | |
|---|---|---|---|---|---|---|---|---|
| | Estimate | 95% CI | Estimate | 95% CI | Estimate | 95% CI | Estimate | 95% CI |
| Conditional $R^2$ | 0.028 | | | | 0.078 | | 0.054 | |
| Marginal $R^2$ | 0.005 | | | | 0.010 | | 0.009 | |
| Model *N* | 2950 | | | | 2557 | | 2557 | |
| Age | 0.0278 | [-0.0113 – 0.0669] | 0.0219 | [-0.0246 – 0.0684] | 0.0087 | [-0.0334 – 0.0509] | 0.0161 | [-0.0260 – 0.0582] |
| $Age^2$ | -0.0057 | [-0.0464 – 0.0351] | -0.0145 | [-0.0635 – 0.0345′ | 0.0103 | [-0.0338 – 0.0543] | 0.0262 | [-0.0178 – 0.0703] |
| Sex [Male] | -0.0128 | [-0.1118 – 0.0862] | -0.0835 | [-0.1997 – 0.0328] | **-0.1354** | **[-0.2408 – -0.0300]** | -0.1248 | [-0.2302 – -0.0194] |
| Pubertal timing | -0.0510 | [-0.1157 – 0.0138] | -0.0054 | [-0.0792 – 0.0685] | 0.0383 | [-0.0302 – 0.1067] | -0.0472 | [-0.1157 – 0.0212] |
| Puberty tempo | -0.0398 | [-0.1062 – 0.0266] | -0.0611 | [-0.1370 – 0.0147] | -0.0025 | [-0.0729 – 0.0680] | 0.0198 | [-0.0507 – 0.0902] |
| Pubertal timing x Sex [Male] | 0.0169 | [-0.0830 – 0.1168] | 0.0321 | [-0.0866 – 0.1509] | -0.0581 | [-0.1652 – 0.0490] | -0.0156 | [-0.1227 – 0.0915] |
| Puberty tempo x Sex [Male] | 0.0666 | [-0.0171 – 0.1502] | 0.0442 | [-0.0531 – 0.1415 ] | -0.0210 | [-0.1103 – 0.0684] | -0.0434 | [-0.1328 – 0.0459] |

*Note*. These report regression coefficients and associated 95% confidence intervals for linear mixed effects models of the general form shown in Eq. 3 . Bold values indicate significant predictors at $p_{FWE} < 0.05$.
